# Bitter Melon (*Momordica Charantia*) Supplementation Has no Effect on Hypercholesterolemia and Atherosclerosis in Mice

**DOI:** 10.1101/2020.04.27.043430

**Authors:** Shayan Mohammadmoradi, Deborah A. Howatt, Hong S. Lu, Alan Daugherty, Sibu P. Saha

**Affiliations:** Saha Cardiovascular Research Center, University of Kentucky, Lexington, KY, USA; Department of Pharmacology and Nutritional Sciences, University of Kentucky, Lexington, KY, USA; Department of Physiology, University of Kentucky, Lexington, KY, USA; Department of Surgery, University of Kentucky, Lexington, KY, USA

**Keywords:** *Momordica Charantia*, atherosclerosis, hypercholesterolemia, Cholesterol

## Abstract

*Momordica Charantia,* commonly known as bitter melon, has been reported to ameliorate diet-induced obesity and dyslipidemia. However, the effects of *M. Charantia* on atherosclerosis have not been determined. This study investigated the effects of *M. Charantia* on diet-induced atherosclerosis in LDL receptor deficient mice. Female mice (6-8 weeks old) were fed a saturated fat-enriched diet. In group 1, mice were fed this diet alone, while mice in groups 2 and 3 were fed the diet supplemented with *M. Charantia* either 0.1% or 1% by weight, for 12 weeks. No significant differences in body weights were observed among the 3 groups. There were also no significant differences in plasma cholesterol concentrations among the 3 groups. To determine the effects on atherosclerosis, lesion areas were measured on aortic intima by an *en face* technique. Neither dose of *M. Charantia* supplementation changed atherosclerosis development.

## 1. Introduction

Atherosclerosis is one of the major cardiovascular diseases with high morbidity and mortality rates in different populations with hypercholesterolemia and obesity as major risk factors.(1,2) Lifestyle modification and healthy dietary habits reduce the risks of predisposition to atherosclerosis and overall cardiovascular health.(3) However, successful lifestyle changes are challenging. Complementary approaches and dietary adjuncts that identify potential therapeutic targets on hypercholesterolemic properties have substantial interests.

*Momordica Charantia,* colloquially known as bitter melon, is an herbal product that has been used traditionally for treatment of diabetes.(4) It has been reported in both human and animal studies that *M. Charantia* has beneficial effects on adiposity and lipid metabolism.(5–8) In one study, *M. Charantia* supplementation reduced plasma cholesterol and triglyceride concentrations in normolipidemic rats. Moreover, aqueous extracts of *M. Charantia* fruit improved lipid profile in normolipidemic diabetic rats. Although beneficial effects of *M. Charantia* on lipid metabolism in normolipidemic state have been demonstrated, there is a lack of information on anti-atherosclerosis effect in hypercholesterolemic animals.

Low-density lipoprotein receptor *(Ldlr)* deficient mice are a commonly used animal model to study atherosclerosis. (1) These mice have profoundly increased plasma cholesterol concentrations when fed a saturated fat-enriched diet. In a quest to determine the role of *M. Charantia* supplementation on atherogenesis, the present study was performed to determine whether modest or substantial dietary supplementation of *M. Charantia* would attenuate atherosclerosis development in *Ldlr^-/-^* mice.

## 2. Materials and methods

### 2.1. Mice and diet

Female *Ldlr^-/-^* mice (B6.129S7-Ldlrt^m1Her^/J; stock #002207) were purchased from The Jackson Laboratory (Bar Harbor, ME, USA). Mice were fed a saturated fat-diet (42% kcal/wt from milk fat diet no. TD.88137; Envigo, Madison, WI, USA) in group 1 for 12 weeks. The Western diet was also consumed by groups 2 and 3 with supplementation of a moderate (0.1% wt/wt; TD. 170736) or high (1% wt/wt; TD. 170735) amount of *M. Charantia* extract, respectively (n=10 per group). **Figure 1A** shows the study design. The mice were maintained in a light:dark cycle of 14:10 hours. Mice were 8 weeks old at the initiation of the study. All procedures were approved by the University of Kentucky Institutional Animal Care and Use Committee.

**Figure 1:**
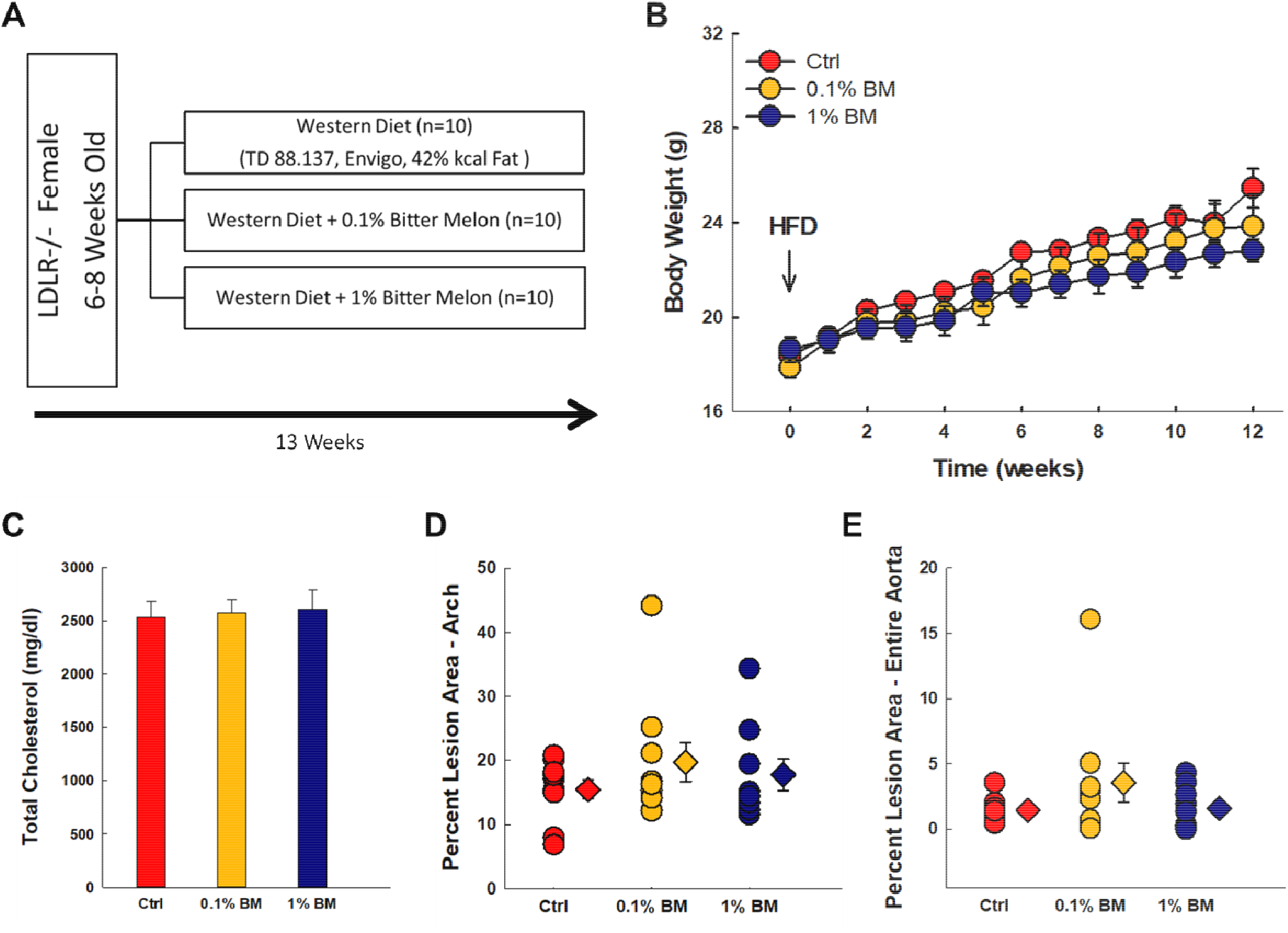
Female LDLR deficient mice were fed a high-fat diet supplemented with *M. Charnatia* for 12 weeks. (A) Body weights were measured weekly. (B) *M. Charantia* supplementation did not show any effect on total plasma cholesterol concentrations. (C) Atherosclerotic lesion development. (D-E) Percent of atherosclerotic lesion areas were measured on the intimal surface of ascending aorta and arch (D) and entire aorta. (E) Diamonds represent group means and error bars are SEMs. Circles represent individual mice. Ctrl; control, BM; Bitter melon. HFD; high fat diet.

### 2.2. Measurement of plasma cholesterol concentrations

Plasma cholesterol concentrations were measured using an enzymatic kit (cat. no. 999-02601; Wako Chemicals USA, Richmond, VA, USA).

### 2.3. Quantification of atherosclerosis

Atherosclerosis was quantified on the intimal surface of the proximal aortic region (ascending aorta, arch, and from the aortic orifice of left subclavian artery to 3 mm below) or the entire aorta from the ascending to the iliac bifurcation by an *en face* method as described previously,(9) and in accord with the guidelines published by the American Heart Association. (1)

### 2.4. Statistical Analyses

SigmaPlot 13.0 (Systat Software Inc., San Jose, CA, USA) was used for all statistical analyses. To compare multiple groups that passed normality and equal variance tests, One Way ANOVA with Holm-Sidak post hoc test was used. Kruskal-Wallis One Way ANOVA on Ranks and Dunn’s post hoc test was used for data that failed normality or equal variance test. P<0.05 was considered statistically significant.

## 3. Results

### 3.1. *M. Charantia* supplementation had no effect on body weights

*M. Charantia* has a bitter taste. Rodents are sensitive to inclusion of extra compounds to their diet, which can consequently affect food intake. To determine the impact on the inclusion of this supplement, mice were weighed weekly during the study period. All groups had steady weight gain that was not significantly different among the 3 groups (**Figure 1B**).

### 3.2. Plasma cholesterol concentrations were not affected by *M. Charantia*

Since hypercholesterolemia has a direct association with atherogenesis, plasma cholesterol concentrations were measured at the end of the study. All 3 groups had concentrations of approximately 2,500 mg/dl that were not significantly different among groups (**Figure 1C**).

### 3.3. *M. Charantia* had no effect on atherosclerosis

Atherosclerosis lesion areas were quantified on the intimal surface of the ascending aorta and arch region, or the entire aorta. All mice had atherosclerotic lesions but there was no significant difference among groups (**Figure 1D-E**)

## 4. Discussion

*M. Charantia* has been shown to promote beneficial effects on lipid metabolism. In this study, we investigated the role of *M. Charantia* supplementation on development of atherosclerosis in hypercholesterolemic mice. We hypothesized that supplementation of *M. Charantia* would have a dose-dependent anti-atherosclerotic effect in female *Ldlr^-/-^* mice. However, 12 weeks of *M. Charantia* supplementation did not have effect on plasma cholesterol concentrations or atherosclerosis development in this study.

*M. Charantia* extracts have lipid lowering effects on both diabetic and high fat diet-fed rats. (6, 8, 10–12) Aqueous extracts of *M. Charantia* not only improved glucose tolerance but also lowered plasma apolipoprotein B (apoB)-100 and apoB-48 concentrations in female C57BL/6 mice fed a high fat diet. (13) A 30-day administration of *M. Charantia* fruit extract also decreased triglyceride and LDL cholesterol concentrations in diabetic rats, (14) and *M. Charantia* (0.75% wt/wt) supplementation reduced plasma cholesterol concentrations in high fat diet fed rats. (8) With many chemicals contained in the extract, *M.CharantiaS* active hypoglycemic and hypolipidemic compounds are unknown. The known isolated compounds such as charantin (a steroid glycoside) and a 166 residue insulin mimetic peptide have been reported to be potential compounds responsible for such beneficial effects. (15)

In agreement with the present study, there are inconclusive results in studies investigating the role of this traditional plant on dyslipidemia and hypercholesterolemia-induced atherosclerosis. Earlier studies reported no changes in either metabolic parameters or lipid metabolism after different doses of *M. Charantia* supplementation. (8, 10, 16) Rats fed diets for 14 days containing *M. Charantia* freeze-dried powder supplemented with 1% wt/wt had no change in plasma cholesterol concentrations. (10) Also, a clinical study using encapsulated *M. Charantia* for a duration of 3 months did not report any significant difference in the lipid profile of patients. (17) Another randomized, double-blind, placebo-controlled trial also reported no significant effect on mean fasting blood sugar, total cholesterol, and weight after treatment with *M. Charantia* capsules.(18)

Although our own studies contrast some of the previously published results, these discrepancies may be attributed to different study designs used in combination with the different preparations, the range of doses, and administration modes of the *M. Charantia.* In addition, many recent reports indicate that treatment and outcomes of atherosclerosis exhibit sex differences in clinical studies and sex differences have been widely reported in mouse atherosclerotic studies. (19) Our studies used female mice which may be another compound factor to the divergent results. (19) Moreover, confirmation of extraction and purification of *M. CharantiaS* active compounds and concentration measurement of such compounds in plasma would be helpful additions for future studies.

In summary, our study results indicate that supplementation of rodent diet with moderate (0.1% wt/wt) or high (1% wt/wt) doses of *M. Charantia* has no effects on hypercholesterolemia and atherosclerosis development. Further studies should investigate the effective dose of supplementation with measuring the bioactive compounds as a marker of consumption.

## References

1. Daugherty A, Tall AR, Daemen MJAP, Falk E, Fisher EA, García-Cardeña G, et al. Recommendation on Design, Execution, and Reporting of Animal Atherosclerosis Studies: A Scientific Statement From the American Heart Association. Arteriosclerosis, Thrombosis, and Vascular Biology. 2017;37(9):e131–e57.

2. Bornfeldt Karin E, Tabas I. Insulin Resistance, Hyperglycemia, and Atherosclerosis. Cell Metabolism. 2011;14(5):575–85.

3. Artinian NT, Fletcher GF, Mozaffarian D, Kris-Etherton P, Van Horn L, Lichtenstein AH, et al. Interventions to Promote Physical Activity and Dietary Lifestyle Changes for Cardiovascular Risk Factor Reduction in Adults. A Scientific Statement From the American Heart Association. Circulation. 2010;122(4):406–41.

4. Alam MA, Uddin R, Subhan N, Rahman MM, Jain P, Reza HM. Beneficial Role of Bitter Melon Supplementation in Obesity and Related Complications in Metabolic Syndrome. Journal of Lipids. 2015;2015.

5. Shih C, Lin C, Lin W. Effects of Momordica Charantia on Insulin Resistance and Visceral Obesity in Mice on High-Fat Diet. Diabetes research and clinical practice. 2008;81(2):134–43.

6. Huang HL, Hong YW, Wong YH, Chen YN, Chyuan JH, Huang CJ, et al. Bitter Melon (Momordica Charantia L.) Inhibits Adipocyte Hypertrophy and Down Regulates Lipogenic Gene Expression in Adipose Tissue of Diet-Induced Obese Rats. British Journal of Nutrition. 2008;99(2):230–9.

7. Chen PH, Chen GC, Yang MF, Hsieh CH, Chuang SH, Yang HL, et al. Bitter Melon Seed Oil–Attenuated Body Fat Accumulation in Diet-Induced Obese Mice Is Associated with cAMP-Dependent Protein Kinase Activation and Cell Death in White Adipose Tissue. The Journal of Nutrition. 2012;142(7):1197–204.

8. Chan LLY, Chen Q, Go AGG, Lam EKY, Li ETS. Reduced Adiposity in Bitter Melon (Momordica charantia)–Fed Rats Is Associated with Increased Lipid Oxidative Enzyme Activities and Uncoupling Protein Expression. The Journal of Nutrition. 2005;135(11):2517–23.

9. Daugherty A, Whitman SC. Quantification of Atherosclerosis in Mice. In: Hofker MH, van Deursen J, editors. Transgenic Mouse: Methods and Protocols. Totowa, NJ: Humana Press; 2002. p. 293–309.

10. Jayasooriya AP, Sakono M, Yukizaki C, Kawano M, Yamamoto K, Fukuda N. Effects of Momordica Charantia Powder on Serum Glucose Levels and Various Lipid Parameters in Rats Fed With Cholesterol-Free and Cholesterol-Enriched Diets. Journal of Ethnopharmacology. 2004;72(1):331–6.

11. Chaturvedi P, George S. Momordica charantia Maintains Normal Glucose Levels and Lipid Profiles and Prevents Oxidative Stress in Diabetic Rats Subjected to Chronic Sucrose Load. Journal of Medicinal Food. 2010;13(3):520–27.

12. Zeng Y, Guan M, Li C, Xu L, Zheng Z, Li J, et al. Bitter Melon (Momordica Charantia) Attenuates Atherosclerosis in Apo-E knock-out Mice Possibly Through Reducing Triglyceride and Anti-Inflammation. Lipids in Health and Disease. 2018;17(1):251.

13. Shih CC, Lin CH, Lin WL. Effects of Momordica charantia on insulin resistance and visceral obesity in mice on high-fat diet. Diabetes research and clinical practice. 2008;81(2):134–43.

14. Chaturvedi P, George S. Momordica charantia Maintains Normal Glucose Levels and Lipid Profiles and Prevents Oxidative Stress in Diabetic Rats Subjected to Chronic Sucrose Load. Journal of Medicinal Food. 2010;13(3):520–7.

15. Nerurkar PV, Lee YK, Motosue M, Adeli K, Nerurkar VR. Momordica Charantia (Bitter Melon) Reduces Plasma Apolipoprotein B-100 and Increases Hepatic Insulin Receptor Substrate and Phosphoinositide-3 Kinase Interactions. British Journal of Nutrition. 2008;100(4):751–9.

16. Chen Q, Chan LYC, Edmund TSL. Bitter Melon (Momordica charantia) Reduces Adiposity, Lowers Serum Insulin and Normalizes Glucose Tolerance in Rats Fed a High Fat Diet. The Journal of Nutrition. 2003;133(4):1088–93.

17. Tsai CH, Chen ECF, Tsay HS, Huang CJ. Wild Bitter Gourd Improves Metabolic Syndrome: A Preliminary Dietary Supplementation Trial. Nutrition Journal. 2012;11:4-.

18. Dans AML, Villarruz MVC, Jimeno CA, Javelosa MAU, Chua J, Bautista R, et al. The effect of Momordica charantia Capsule Preparation on Glycemic Control in Type 2 Diabetes Mellitus Needs Further Studies. Journal of Clinical Epidemiology. 2007;60(6):554–9.

19. Robinet P, Milewicz Dianna M, Cassis Lisa A, Leeper Nicholas J, Lu Hong S, Smith Jonathan D. Consideration of Sex Differences in Design and Reporting of Experimental Arterial Pathology Studies—Statement From ATVB Council. Arteriosclerosis, Thrombosis, and Vascular Biology. 2018;38(2):292–303.

